# Cell state and transcription factor modulation during extended ex vivo CD8^+^ T-cell expansion

**DOI:** 10.1101/2024.07.17.603780

**Authors:** Yuan Lui, Edward Jenkins, Emily Zhi Qing Ng, Mateusz Kotowski, Sydney J Mullin, Joseph Clarke, Simon J Davis, Ana Mafalda Santos, Sumana Sharma

**Author notes:** authors contributed equally (Yuan Lui, Edward Jenkins).

## Abstract

Adoptive cell therapy is becoming a cornerstone of tumour immunotherapy. It relies on the relatively long-term (> 2 week) ex vivo expansion of T cells either in the form of tumour-infiltrating cells, or bulk cells modified with the expression of heterologous signalling proteins, e.g., chimeric antigen receptors. However, relatively little is known about the developmental trajectories of T cells under these conditions at the system level, or whether the pathways governing these trajectories could be manipulated for clinical advantage. Using bulk RNA-seq analysis of T cells expanded and rested over a 17-day period, we produce a resource revealing how gene expression changes as cells transition through distinct cellular states over the course of activation and ex vivo expansion. By integrating this resource with published single-cell RNA-seq data, we identify a member of the AP1 transcription factor (TF) family, FOSL1, that primes CD8^+^ T-cells towards an effector/killing phenotype. Remarkably, FOSL1 over-expression during T-cell expansion produced ‘super engager-like’ T-cells, evidenced by their gene-expression signatures and enhanced cancer-cell killing capacity. This establishes proof-of-principle for the rational engineering of T cells via TF modification during ex vivo expansion, offering a route to improving adoptive T-cell therapy.

## Introduction

Adoptive cell therapy (ACT) with ex vivo-generated T-cells is an emerging approach for treating human malignancies. Despite remarkable successes, there remain significant barriers to the efficacy of ACT as durable responses to ACT are still difficult to achieve in some patients (Restifo et al, 2016; Borcoman et al, 2019). Alongside tumour heterogeneity and microenvironment, the shortcomings of existing cell therapies can be attributed, at least in part, to the cellular state of the initial and expanded T-cells used, which impacts their proliferation and differentiation, as well as their eventual homing, persistence, and cytotoxicity (Fraietta et al, 2018). Optimising the T-cell state during the manufacturing process is therefore key to improving treatment efficacy.

Current approaches for effecting changes in the T-cell state for ACT utilise different starting subpopulations or chemical and genetic modification during ex vivo expansion (Rohaan et al, 2019). It is thought that less well-differentiated T-cell subsets (i.e., naïve, central memory, or memory stem cells) have functional advantages and are considered a better source of starting material than more differentiated cells (Hinrichs et al, 2011; Klebanoff et al, 2012; Xu et al, 2014; Busch et al, 2016). Less differentiated cells, however, are harder to manipulate genetically, and so protocols have been developed that utilise chemical inhibition of signalling pathways (e.g., p38 kinase or AKT signalling) to allow efficient transduction and expansion of minimally differentiated cells (Klebanoff et al, 2017). T-cell optimization has also been sought by expressing chimeric antigen receptors (CARs) with tailored specificity and by altering downstream signalling through the knockout (KO)/loss of function of proteins such as PD-1, RASA2, PTPN2, SOCS1, and through gains of function using CRISPRa (Gurusamy et al, 2020). As a cell’s state is determined by the activity of transcription factors (TF), modifying TF expression in T cells during their ex vivo manufacture could be an alternative approach to improving T-cell quality and for tailoring ACT. Indeed, it was recently shown that over-expressing the TF AP4 in CAR T-cells led to enhanced fitness of chronically stimulated CAR T-cells. Whilst this approach is promising, it has not been possible to use it systematically beyond performing technically demanding over-expression screens (Blaeschke et al, 2023). An alternative approach would be to use in silico modelling to predict the appropriate TF target modification needed to produce a desired cell state. However, this would require a high-resolution temporal map revealing how gene-expression networks change over the course of T-cell expansion ex vivo, as well as an understanding of how TF networks establish the represented cell state and of causality between TF networks and T-cell state trajectories, a resource that is currently lacking.

Owing to advances in single-cell sequencing, we now know that T cells exist in a variety of states bounded by well-defined axes depending on the signals received (Papalexi & Satija, 2018). Existing approaches to characterizing cell state rely on using techniques such as single-cell RNA-seq and CyTOF/spectral flow cytometry. However, the former typically samples a subset of RNA (i.e., generates a high ‘dropout’ rate or genes with count values of zero, while the latter only samples up to ∼40 proteins. While newer approaches are being developed to infer transcriptional signatures using single-cell transcriptomic datasets, owing to the low expression of TFs these approaches may not always be helpful (Hicks et al, 2018; Bacher & Kendziorski, 2016; Müller-Dott et al, 2023). More recent developments have relied on in silico perturbation experiments to identify TFs that control cell state, e.g., CellOracle (Kamimoto et al, 2023), and while powerful inference tools, these still rely on (incomplete) single-cell RNA-seq data sets to make such inferences. Furthermore, single-cell RNA-seq experiments typically focus on a few time points mostly due to the high costs associated with the method, reducing the temporal resolution of TF analyses. As in silico perturbation tools can only operate within measured state space, limited temporal datasets also reduce the power of these approaches. Although a high temporal-resolution, single-cell RNA-seq analysis of T-cell activation up to 5 days has been undertaken (Soskic et al, 2022), this did not include a period of ‘rest’ that T cells prepared for ACT are typically given after an initial round of activation, which is known to restore functionality and induce epigenetic reprogramming in, e.g., exhausted human CAR-T cell populations (Weber et al, 2021). There remains, therefore, the need for a complete analysis of TF expression across the whole of the ex vivo expansion. Compared to single-cell RNA-seq, bulk RNA-seq can allow for much deeper dissections of TF networks over time for populations that are homogenous, which is achievable ex vivo.

Cells generated for ACT are produced ex vivo using reagents that engage the T-cell receptor (TCR) and the co-stimulatory receptor CD28 (e.g., by using beads coated with anti-CD3 and anti-CD28 antibodies). This leads to the rapid expansion of T cells over two weeks, essential for producing the number of cells required for therapy. While cell expansion is key for all forms of ACT, the extent of malleability of T cells during T-cell expansions under conditions widely used for ACT is not fully explored. Here, we use bulk RNA-seq to generate a map of gene expression trajectories derived from ex vivo activated, expanded, and rested naïve CD8^+^ T-cells across 17 days. We integrated the data with a previously published catalog of T-cell developmental stages to categorise which of these stages are best represented by ex vivo expanded cells. We then use this resource to categorise key TF networks that define different states created during the T-cell activation process. In addition, we use CellOracle (Kamimoto et al, 2023) to perform in silico perturbations of TFs and analyse their impact on cell state transitions. By integrating our bulk RNA-seq time course data with predictions from CellOracle, we identify TF targets that are predicted to drive the core behaviours of T cells during the early stages of T-cell activation. As proof of principle for our approach, we establish FOSL1, an AP1 family protein, as a new promoter of an enhanced effector state in CD8^+^ T-cells, characterised by increased cytokine production, the gene signatures of ‘super-effectors’, killing, and the expression of tissue-homing receptors.

## Results

### T-cell states and transitions during 17-day ex vivo activation and expansion

To achieve a comprehensive overview of cell states and TF network changes during ex vivo T-cell activation, we performed bulk RNA-seq on stimulated primary CD8^+^ T-cells starting from a naïve population over the course of 17 days. This period was chosen to reflect the activation timeline that T cells typically experience during infections in vivo (Kaech & Cui, 2012; Araki et al, 2017), and expansion ex vivo for ACT (Watanabe et al, 2022). We used naïve CD8^+^ T cells as a starting population to reduce heterogeneity for our bulk RNA-seq measurements and because, versus unfractionated T-cells, these cells exhibit a superior capacity to expand, persist and eliminate tumours (Hinrichs et al, 2011). The naïve CD8^+^ T-cells were stimulated with anti-CD3/CD28 beads for 7 days, and RNA was extracted at defined time-points during the activation stage (i.e., 30 mins, 3 h, 12 h, 24 h, 48 h, 72 h, and 7 days). The beads were then removed, and RNA extracted following 2-, 7-, or 10-days of rest. For each time-point we extracted total RNA and performed bulk RNA-seq in three technical replicates. Initial principal component analysis (PCA)-based examination of the activation time-course revealed six major clusters of cells distributed across the stimulation timepoints (Fig. 1A). The naïve state and cells 30 minutes post-activation grouped together but by 3 hours the T-cell state diverged substantially from that of the naïve cells. The next set of four time points (12-72 h) grouped together but remained distinct from T cells activated for 3 h. Cells from the 7-day time point, and 7-days activation with 2-days rest, diverged from each other and other stimulated conditions. Lastly, cells from the long-rest timepoints (7-or 10-days rest) grouped together.

**Figure 1:**
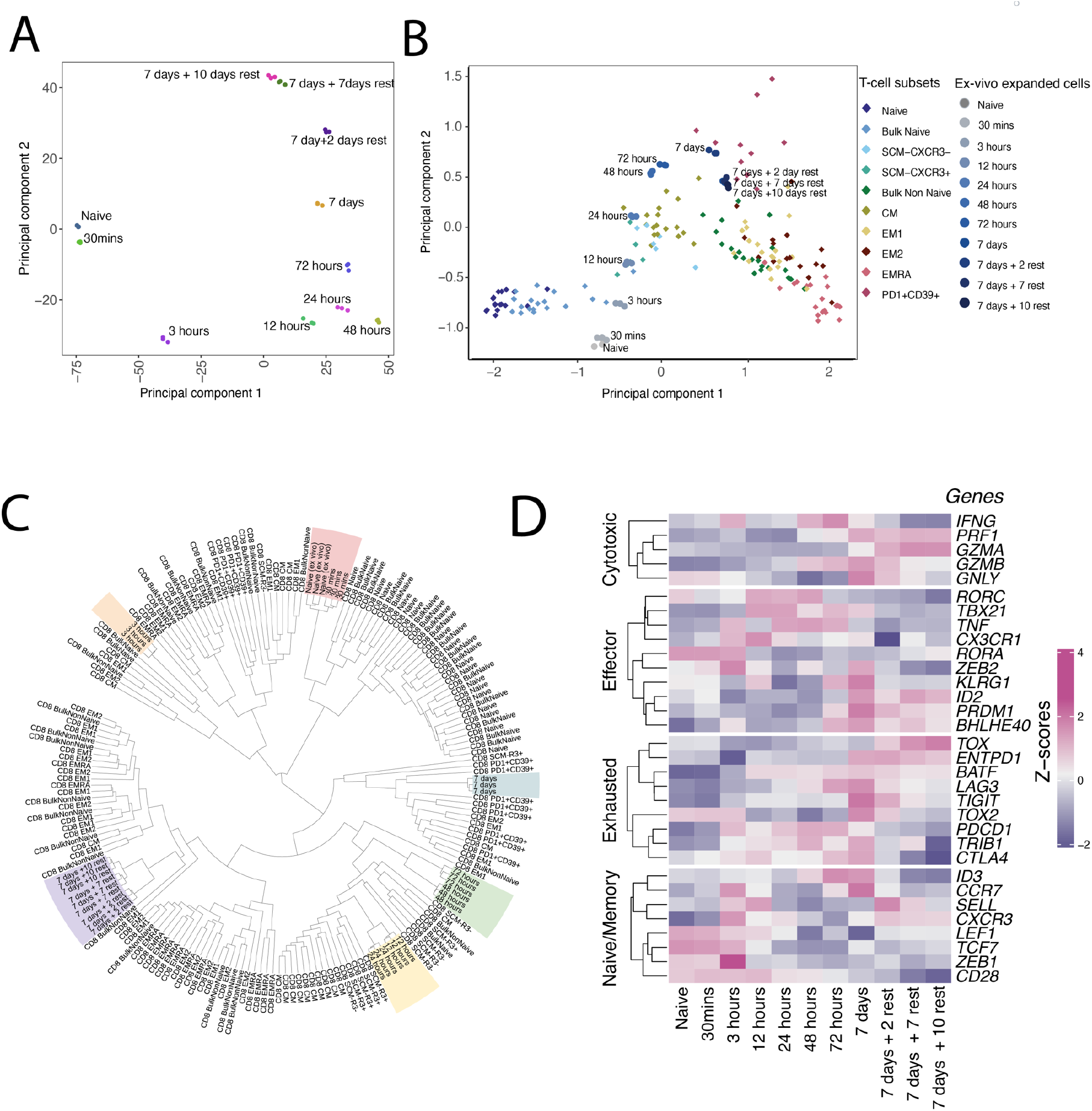
In vitro stimulated CD8+ T cells recapitulate physiological T-cell states. A. PCA of naïve CD8 T cells activated using CD3/CD28 beads over the course of 17 days. B. PCA post batch correction of catalogue of healthy T-cell subsets integrated with ex vivo stimulated cells. C. Distance matrix-based hierarchical clustering of the integrated dataset. D. Heatmap of the expression of marker genes defining different T-cell subsets.

Next, we compared the transcriptomic profiles of the in vitro-expanded CD8^+^ cells with those of T-cell subsets isolated from the blood of healthy human donors analysed using RNA-seq and catalogued by Giles et al. (Giles et al, 2022). This catalogue comprises, naïve, stem-cell memory (SCM), central memory (CM), effector memory (EM), CD45RA^+^ effector memory (EMRA), and putative exhausted (Tex) CD8^+^ T-cells. SCM cells were further divided into two groups based on the expression of CXCR3 (SCM-CXCR3^-^ and SCM CXCR3^+^), and EM cells divided into two groups based on the expression of CD27 (EM1 (CD27^+^) and EM2 (CD27^-^)). As an initial examination, we reprocessed the published dataset and integrated the two datasets post batch-correction with ComBat (Suppl. Fig. 1) (Zhang et al, 2020). This allowed direct comparison of the gene expression profiles of ex vivo-expanded cells with those of natural CD8^+^ T-cells. An initial PCA examination of the integrated dataset arrayed the pre-defined subsets along PC1 with naïve CD8^+^ T-cells located at one end, memory subsets (SCM and CM) in the middle, and EMRA CD8^+^ T-cells at the opposite end (Fig. 1B). The ex vivo expanded cells, on the other hand, were distributed along the PC2 axis with naïve cells at the bottom and cells stimulated for seven consecutive days on the other end. The overlay of the two samples showed the distinct T-cell states that each ex vivo expanded T-cell represented. We additionally used distance-matrix clustering to assess the similarity between the expanded cells and the different T-cell subsets (Fig. 1C). Naïve and 30-minute cells were clustered together close to the naïve cells from the catalogue of T-cell subsets. Within three hours, the cells were transiting towards the SCM cell population but were closest to the bulk naïve cell population. Cells stimulated for 12-24 hours were most like the SCM subset. Further expansion of the cells produced cells more like the CM population and eventually similar to the PD-1-expressing putative exhausted state. Interestingly, rested cells post 7-day stimulation were distinct from the 7-day stimulated but unrested cells. Rested cells were more like the CM and EM1 populations. No ex vivo-expanded cell populations were similar to the EMRA population.

To further validate the T-cell states, we examined the key cell-state markers previously defined by Giles et al. and found that the activated cells expressed effector markers including *TBX21, RORC*, and *TNF* rapidly upon stimulation followed by markers of cytotoxicity, e.g., genes encoding granzymes and perforins including *GZMB, GZMA*, and *PRF1*, during the later stages as stimulation proceeded. Naïve and memory markers were either expressed early or late during the stimulation time course. Genes encoding several inhibitory receptors, i.e., PD-1, LAG-3, CTLA-4, and TIGIT were expressed on Day 7 but their expression was downregulated following the removal of stimulation, consistent with the rested cells being less putatively exhausted (Fig. 1D). Altogether, this analysis showed that in vitro activation drives the formation of a succession of physiological CD8^+^ T-cell states each of which is, in principle, selectable.

### Gene expression profiles and patterns during 17-day ex vivo activation and expansion

To further characterise the different states of ex vivo-expanded T-cells, we next examined individual genes and enriched pathways at the different stages of activation. We identified differentially expressed genes at each time-point compared to the naïve state and used pathway enrichment to identify how the pathways changed over the course of the activation period. Gene pathway analysis showed that at 3 h of stimulation, signalling components (e.g., TCR signalling, MAPK signalling, JAK-STAT signalling, Hippo signalling, NF-kappa B signalling) were significantly upregulated (Fig. 2A). Around the peak of activation at 12-72 h, the cells entered a highly proliferative state (or expansion phase) with enrichment of pathways relating to DNA replication, ribosome biosynthesis, proteasomes, cell cycle, and metabolism. During this period, pathways relating to apoptosis and cellular senescence were also enriched, likely priming the T cells for the contraction phases normally observed after pathogen clearance. Strikingly, although genes associated with the cell cycle and DNA replication persisted through the entire time course (i.e., even after the activation stimulus was removed), pathways relating to ribosomal biogenesis were no longer enriched at the 7-day activation and rest stages. This indicated that by Day 7 the cells had transitioned to a low translational and proliferative state, as observed for T cells in an 8-day mouse-infection model (Araki et al, 2017). During the rest stage, the cells likely became quiescent, as suggested by the reduced enrichment of genes linked to proteasome expression and function.

**Figure 2:**
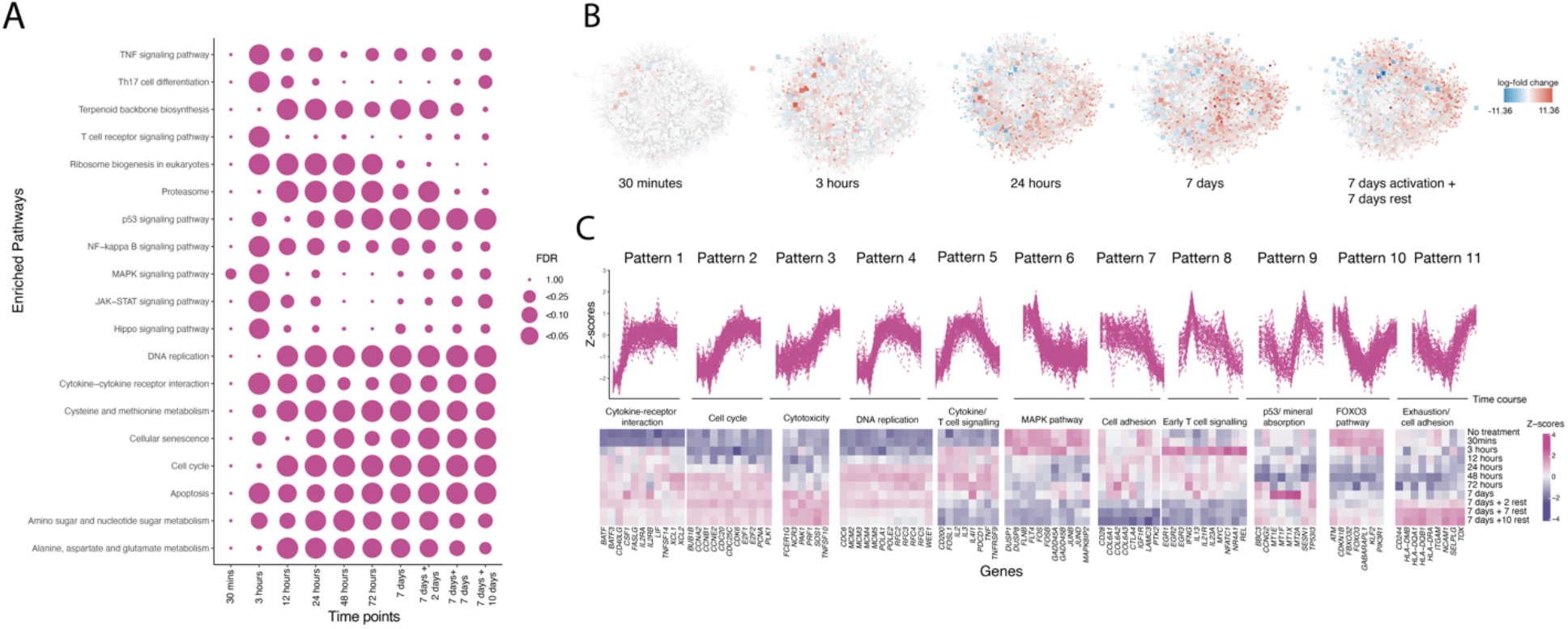
Gene expression trajectories of CD8+ T-cells over the course of 17 days. A. Gene regulatory network of CD8+ T-cells generated using GENIE3. Fold change values compared to naïve state are overlaid on the network. B. Pathway enrichment for genes enriched at each time-point compared to the naïve state. C. 11 distinct gene expression patterns and representative genes in key pathways enriched in the time course dataset.

To further identify distinct regulatory networks during the activation time course, we then examined changes in the gene-expression networks. We extracted the 35% most variably expressed genes across all timepoints and generated a gene-regulatory network using GENIE3 (Huynh-Thu et al, 2010). The resulting network consisted of ∼1600 nodes and ∼7000 edges (complete network in Suppl. Table 1). We then overlaid the fold-change values from the differential gene expression analysis on the nodes to identify regions of the network that were “hot” (indicating increased expression) and “cold” (indicating decreased expression) compared to the naïve T-cell state. Consistent with the PCA analysis, the network signatures could be roughly divided into five states: naïve (0-30 mins), early activation (3 h), mid activation (12-72 h), late activation (7 days), and post-rest (7-10 days rest; Fig. 2B). This confirmed that CD8^+^ T-cells transition through multiple states characterised by distinct gene regulatory networks, during ex vivo expansion.

To generate a comprehensive view of the gene-expression trajectories, we used all the 35% most regulated genes across the different timepoints and clustered them according to their temporal expression patterns. Eleven clusters, referred to hereafter as ‘patterns’, emerged from this analysis (Fig. 2C; see Suppl. Table 2 for the identity of each gene in each pattern and Suppl. Fig. 2 pathways enriched in each pattern). Patterns 1, 2, and 3 were defined by low expression in the naïve state, increased expression upon activation, and sustained expression on resting. The timing of the initial increase in expression varied, with genes in Pattern 1 expressed early in activation (12 h onwards), Pattern 2 around 24-48 h and Pattern 3 as late as 72 h. Each pattern was also associated with the enrichment of distinct pathways, Pattern 1 being enriched in cytokine-cytokine receptor interactions with classical T-cell activation markers such as *TNFSF14, CD40LG, IL2RA, IL2RB, XCL1, XCL2* falling into this pattern. Pattern 2 was highly enriched in cell cycle pathway genes (EF1, EF2, CDK family genes) and Pattern 3 was enriched in genes associated with cytotoxicity (*SOS1, NCR3, FAS, PRF1*) (Kelner et al 1994, Kurachi et al, 2014). Pattern 4 was like Pattern 2 except for a slight drop in expression at the rest stage. The main pathway enriched in this pattern was DNA replication (POL family genes, MCM family genes). Pattern 5 was defined by high expression upon activation followed by low expression immediately following the removal of the stimulus on Day 7. Genes linked to inhibitory receptors, such as *PDCD1* and *CD200* were included in this set. We also noted that IL-2, a key cytokine produced by T cells upon activation, was identified in this pattern together with genes encoding several other IL family proteins such as IL-3, IL-13, IL-21R, IL-4l1. Patterns 6 and 7 comprised high expression in the naïve state and decreased expression as stimulation continued and upon resting. Several AP1 transcription factor family members had this pattern, including *JUND, JUNB, FOS, FOSL2*, and *FOSB* (Karin et al, 1997). The most striking change in TFs in the first 3 h was that of the members of Pattern 8, which included the members of EGR family, NR4A family, *NFATC1*, and *RELA*, which are known to be crucial during initial signalling. Pattern 9 on the other hand was defined by peak expression at the point of removal of beads; components of the p53 signalling pathway were enriched in this pattern (*CDKN1A, TP53I3, SESN1, CCNG2, BBC3*). Two patterns exhibiting very different dynamics were Patterns 10 and 11, which had high expression during the early time-points followed by decreased expression upon stimulation and reversion to high expression during the rest phase. Pattern 10 was enriched in the FOXO signalling pathway, and Pattern 11 several classical exhaustion markers such as *TOX* and *CD244* (2B4) but also pathways controlling cell adhesion (*ITGAM, NCAM1, SELPLG*) (Sekine et al, 2020). Overall, this analysis shows that T-cell phenotypes change dramatically during expansions ex vivo.

### In silico perturbations using CellOracle identify early transcriptional regulators of T-cell activation

The bulk transcriptomic data allowed categorization of different TF families and their functions based on their expression patterns and up/down regulation relative to a previous T-cell state. Next, we undertook a case-study analysis of whether expression data from distinct datasets could be integrated and used to link TFs to given T-cell states and function. For this, we used a scRNA-seq dataset recently published by Bibby et al. In that study, human naïve and memory CD4^+^ and CD8^+^ T-cells were first isolated by staining for CD45RA or CD45RO expression and then either left unstimulated or stimulated for 12 or 24 h with anti-CD3 and anti-CD28 antibodies (Bibby et al, 2022). Although this data only covered the first 24 h of activation, because the same method of activating the T cells was used, we could test whether expression patterns from bulk analysis could be integrated with single-cell data.

Initial unsupervised clustering from the single-cell data revealed a gradient of states from stimulated to activated and proliferative, as the cells were subjected to stimulation with the beads, for both the naïve and memory cell population (Fig. 3A). The existence of this dataset, in which naïve cells or unstimulated memory cells transition into proliferating cells, allowed us to investigate the transcriptional regulation of T-cell state transitions using the regulatory network inference tool, CellOracle (Kamimoto et al, 2023). CellOracle models context-specific gene regulatory networks (GRNs) from scRNA-seq data and information about the ‘accessibility’ of genomic regions. We used a pre-curated TF-binding network together with the scRNA-seq data to infer a T-cell activation gene-regulatory network. Once the GRN is inferred, CellOracle is used to simulate global downstream shifts in gene expression post-perturbation. In silico perturbation in CellOracle is visualised as a vector map on the 2D trajectory space in which perturbation scores (PS) are superimposed. A negative PS implies that TF KO delays or blocks transition to an activated/proliferative state whereas a positive PS implies that the loss of TF function promotes differentiation (the 2D trajectory space is shown in the right panel of Fig. 3A; an example of perturbation of NFKB1 in memory cells in Suppl. Fig. 3).

**Figure 3:**
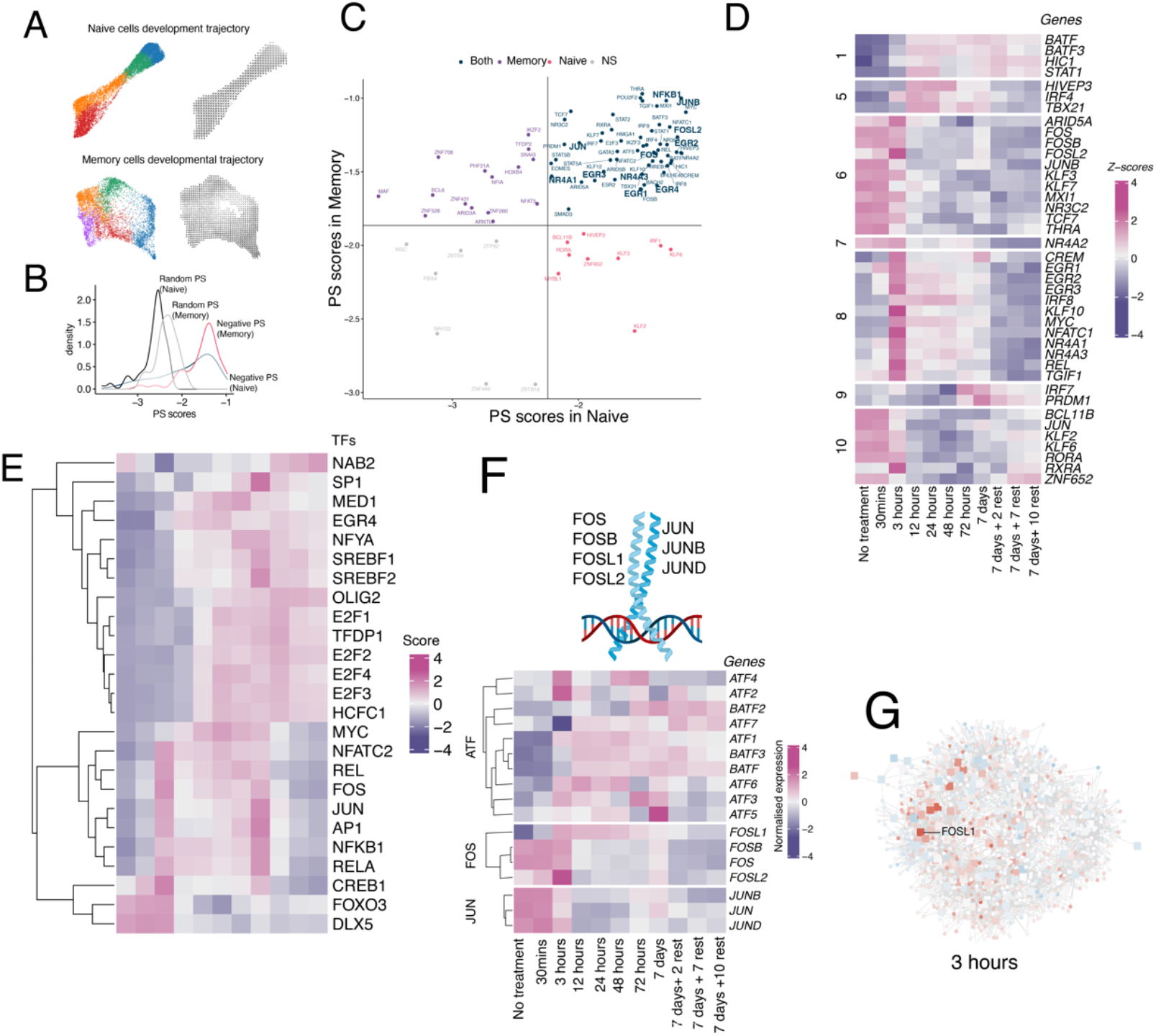
CellOracle predicts the key transcription factors important in early TCR signalling. A. Developmental trajectory of naïve and memory T cells stimulated with anti-CD3/anti-CD28 beads. B. PS score distribution of TFs in memory and naïve CD8+ T cells perturbed using in-silico modeling using CellOracle. C. PS-scores for TFs important for regulation of activation of memory and naïve T cells. D. Dynamic expression patterns as reported by bulk RNA-seq for genes identified from CellOracle as being important for T-cell development for both memory and naïve T-cell development E. Active TFs predicted by CollecTRI during the different stages of ex vivo expansion of T cells. F. Schematic of JUN/FOS family members (top panel). Dynamic expression patterns as reported by bulk RNA-seq for genes in the AP1 family (bottom panel). G. Gene regulatory network of CD8+ T-cells generated using GENIE3. Fold change values at 3 hours compared to naïve state are overlaid on the network.

To validate the use of CellOracle for inferring TF roles in T cells we first identified a set of 132 and 119 transcription factors whose expression was differentially expressed in the three timepoints for the naïve and memory scRNA-seq datasets, respectively. Of these, 88 transcription factors were shared between the two sets. For the overlapping TFs, we systematically performed in silico KOs in the base network and calculated the summed negative perturbation scores for naïve- and memory-cell differentiation over the course of stimulation. We then calculated the negative PS sum cut-off by first calculating PS sum scores from randomised simulations and then setting the cutoff at the 99th percentile to give a cut-off at false-positive rate of 0.01 (score distributions are shown in Fig. 3B). The PS scores were then represented as a scatter plot (Fig. 3C). The resulting distribution of TFs, based on negative PS-scores, was consistent with the known functions of many well-characterised T-cell specific TFs. For example, transcription factors that are known to be involved in the activation and proliferation of T cells, i.e., the AP1 (JUNB, FOSL2, JUN, FOSB, BATF), IRF (IRF4, IRF8), and early growth response (ERG1, ERG2, ERG3), NFKB1, NFATC2, NR4 (NR4A1, NR4A3) families, were located on the top right corner indicating that KO delays early-stage differentiation in both memory and naïve cell population post stimulation.

We then investigated how the expression of the genes predicted to be important for early signalling in T cells according to CellOracle, related to the bulk RNA-seq transcriptional trajectories in the dataset we generated (Fig. 3D). Importantly, 86/88 TFs in this list showed dynamic expression in the first 24 h of stimulation (only IRF7 and PRDM1 from cluster 1 did not). Focusing on Pattern 8 (i.e., genes whose expression increases acutely at 3 h stimulation), 12/15 TFs were also identified with significant PS scores in the CellOracle analysis, confirming these genes as ‘first responders’ during T-cell activation. Genes whose expression patterns were similar in the first 24 hours produced different patterns over the course of the experiment (for example, Pattern 6 vs Pattern 10, and Pattern 1 vs Pattern 5), indicating that it is important to study temporal gene expression during all stages of expansion and not only during initial activation.

We then used our bulk RNA-seq measurements comprising the temporal gene expression changes across the different stages of ex vivo expansion to infer transcription factor activity at different time points using CollecTRI (Müller-Dott et al, 2023). CollecTRI-derived regulons contain signed TF-target gene interactions compiled from 12 different resources to predict transcription factor activity from bulk RNA-seq data (Fig. 3E). This analysis showed that TF activity is markedly different depending on which activation time point the cells have reached. During the early stages, the initial responder TFs are AP1 and NFKB family TFs, whose expression peaks at ∼3h, after which proliferation-related TFs are expressed (E2F family members). AP1 and NFKB family members are expressed up to the point where the stimulus is present (Day 7) but as soon as the stimulus is removed the expression of all initial responder TFs declines.

Both analyses implied that AP-1 is tightly regulated during the ex vivo expansion of T cells. The AP-1 TF complexes are comprised of dimers of JUN (c-Jun, JunB, JunD), FOS (c-Fos, FosB, Fra-1, and Fra-2), ATF (Activating Transcription Factor) (ATF2, ATF3/LRF1, B-ATF), and MAF (musculoaponeurotic fibrosarcoma)(c-MAF, MAFA, MAFB, MAFG/F/K, NRL) proteins (Fig. 3F). The AP1 family is among the most widely studied T-cell activity-related TFs, and the regulation of cell fate by AP-1 is a complex process governed, e.g., by the relative abundances of AP-1 subunits, dimer composition, the nature of stimulation, cell type, and cell environment (Shaulian & Karin, 2002). In T-cells, AP1 TFs comprise both activating-(JUNC, JUND) and exhaustion-promoting factors (JUNB, BATF, BATF3) but the detailed interplay between each of the different AP1 elements in each T-cell state is still to be fully worked out. We examined, especially, the expression of the JUN-, FOS-, and ATF-family members in our bulk RNA-seq data (Fig. 3F). Expression of the FOS, FOSB, and FOSL2 genes was elevated at the naïve stage, but dropped as the cells proliferated. Expression of FOSL1, however, exhibited the opposite behaviour insofar as it was very weakly expressed in naïve cells and increased upon activation, after which it gradually decreased over time. Given the interesting pattern of FOSL1 expression, and because our gene-regulatory network implied that it is one of the key genes expressed at the key 3 h time point (Fig. 3G), and because it is understudied, we examined the role of FOSL1 in T-cell activation.

### Over-expression of FOSL1 in primary CD8^+^ T-cells enhances effector function

To investigate the role of FOSL1 in T-cells, we first generated FOSL1 over-expression and FOSL1 KO lines in immortalised human Jurkat T-cells using lentiviral transduction. These engineered lines were stimulated in activation assays using T-cell stimulator cells (TCS-CD86) that co-express membrane-bound anti-CD3-scFv (to engage the TCR) and high levels of CD86 (to engage the co-stimulatory molecule CD28). This allows signalling strengths to be modulated by pre-blocking the cells with anti-CD86 antibody, or not (see schematics in Fig. 4A). The activation level of Jurkat T-cells was measured via the expression of CD69, a commonly used T-cell activation marker. Jurkat T-cells over-expressing FOSL1 produced high levels of CD69 expression upon stimulation under both high (TCR and CD28 engagement) or low (TCR engagement only) stimulation conditions, when compared to the parental Jurkat T-cell lines (Fig. 4B). Targeting FOSL1 in Jurkat T-cells did not have any impact on the expression of CD69 (Suppl. Fig. 4). This suggested that FOSL1, when expressed at sufficient levels, could act as a positive regulator of T-cell signalling.

**Figure 4:**
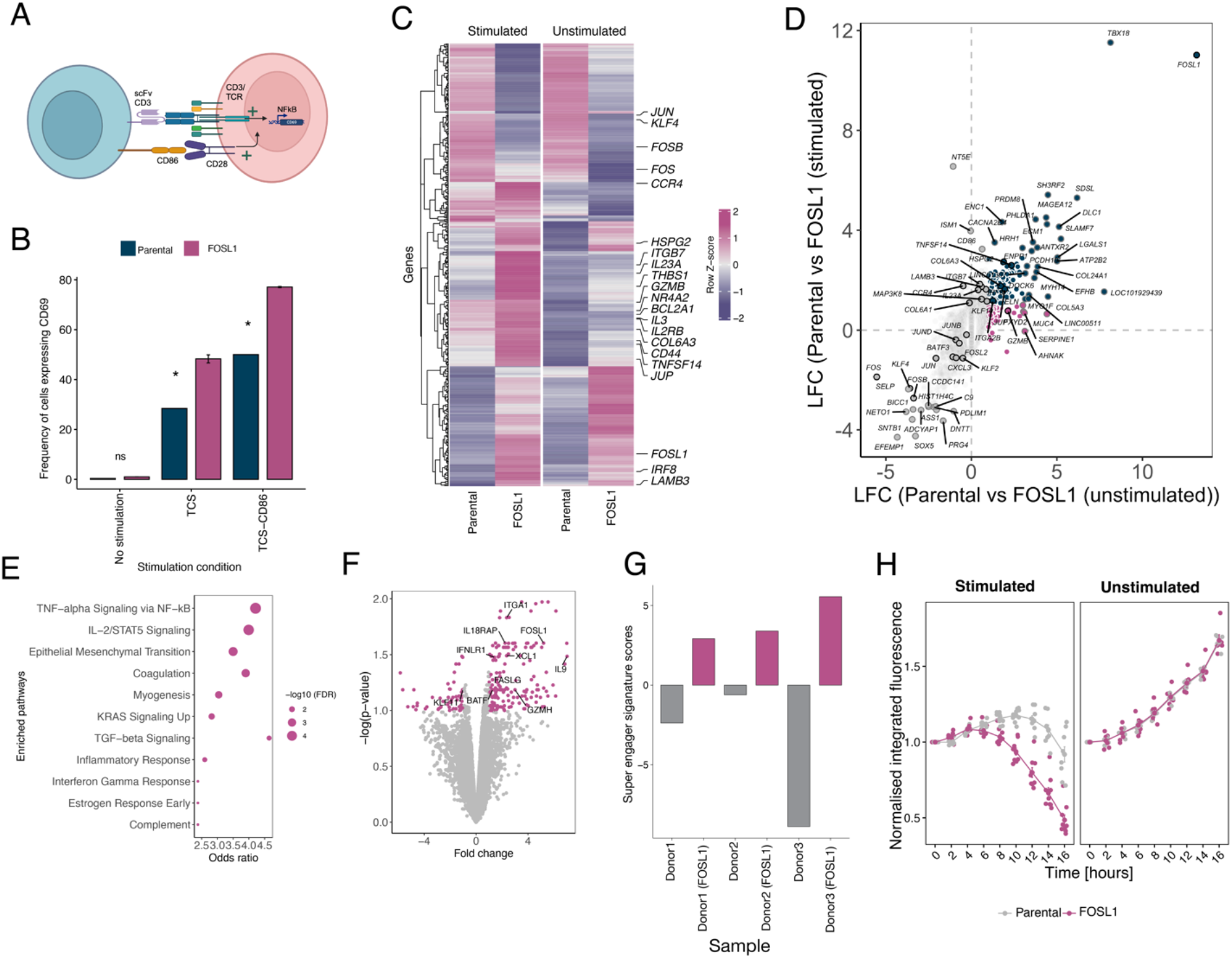
FOSL1 regulates CD8+ T-cell effector function. (A) Schematics of two-cell assay used to study the activation of Jurkat T-cells. (B) Frequency of Jurkat cells expressing CD69 post coculture with stimulator cells. TCS and TCS-CD86 are used as stimulator cells. (C) Z-scores from RNA-seq of genes that are identified as differentially expressed between the comparisons made for cells over-expressing FOSL1 vs matched parental controls with or without stimulation. Key genes are marked. (D) Log-fold change comparisons between cells over-expressing FOSL1 and parental controls in stimulation or non-stimulation conditions. (E) Pathway enrichment for genes over-expressed in FOSL1 over-expressing cell lines compared to parental cell lines under stimulation condition. (F) Volcano plot depicting differentially expressed genes in primary CD8+ T cells over-expressing FOSL1 compared to matched donors. (G) Signatures scores derived from 61 previously classified genes corresponding to ‘super-engager’ phenotype for parental and FOSL1 over-expressing cells. (H) Killing assay for primary CD8+ T cells. Stimulated and unstimulated refers to the conditions in which the primary T-cells were either loaded with the bispecific antibody or left unloaded. The loss of integrated fluorescence indicates killing of target cells (A375) expressing mOrange.

To gain a better insight into the global transcriptional changes elicited by over-expressing FOSL1 in Jurkat T-cells, we performed a bulk RNA-analysis of the parental and FOSL1-over-expressing Jurkat T-cells that were either left unstimulated or were stimulated via the TCR only (i.e., by CD86-blocked TCS-CD86 cells). Interestingly, the transcriptomic profile of the cells over-expressing FOSL1 was already distinct from the parental cell lines prior to stimulation (Fig. 4C). Pathway enrichment suggested that the main differences involved the expression of extracellular matrix organisation, with expression of laminins (LAMA3/5), collagens (COL5A3, COL6A3, COL24A1), and integrins (ITGA3, ITGA2B, ITGAX, ITGAM) enhanced in the FOSL1 expressing lines. Interestingly, several components of AP1 transcription factors themselves, including ATF3, JUN, and JUNB, were downregulated in the FOSL1 over-expressing cell lines. In addition, transcripts encoding granzyme B (GZMB1), a major effector of cytotoxic T-cells, was upregulated in the unstimulated cells. We next compared the difference between parental and FOSL1 over-expressing lines under both stimulated and unstimulated conditions (Fig. 4D). Following stimulation, the key pathways that were enriched upon over-expression of FOSL1 were the IL-2/STAT5 signalling pathway, TNF-alpha signalling via NFKB, and pathways relating to extracellular matrix reorganisation (Fig. 4E). We examined the IL-2 signalling and NFKB signalling pathway in more detail owing to their importance to T-cell effector biology and observed that transcripts for genes encoding regulators of T-cell activation, such as the early activation transcription factor (NR4A2) and cytokines (IL3, IL2RB, IL23A), were enriched in cells expressing FOSL1. Of note, TNFSF14, a major cytokine important for T-cell proliferation, which was predicted to be regulated by FOSL1 in our gene-regulatory network, was upregulated in the FOSL1 over-expressing line. This provided additional evidence that over-expression of FOSL1 enhances signalling in T-cells.

The RNA-seq experiment on Jurkat T-cells suggested that cells over-expressing FOSL1 had increased gene expression of cytokines and granzymes. We then tested if the transcriptional signatures of increased activation would translate to an increase in effector activity in primary cytotoxic CD8^+^ T-cells. For this we transduced three donor CD8^+^ T-cells with lentivirus encoding FOSL1 and performed bulk RNA-seq to assess the transcriptional changes in the cells over-expressing FOSL1 compared to the parental donors. In a similar manner to Jurkat T-cells, the expression of genes relating to the IL-2 pathway were upregulated in FOSL1 over-expressing cells. Effector-specific markers such as *BATF, GZMH*, and *XCL1* were also upregulated suggesting that FOSL1 could have a direct role in CD8^+^ T-cell effector function (Fig. 4F). Strikingly, FOSL1-over-expressing cells were also enriched in previously defined gene signatures of serial-killing super-engagers (Dekkers et al, 2023) (Fig. 4G). FOSL1 over-expressing cells also expressed more Fas ligand (FASLG), which is a cell surface receptor expressed on cytotoxic T-cells that binds Fas receptor on target cells, inducing programmed cell death.

To test if enrichment of the “super engager” transcriptomic signature confers better killing of target cells, we performed target killing assays with parental and FOSL1 over-expressing donor CD8^+^ T-cells. In the killing assay, we first pre-incubated either parental or FOSL1 over-expressing CD8^+^ T-cells with a bispecific molecule, which is an engineered biologic with a melanoma antigen (GP100)-specific TCR fused to a single chain antibody fragment that binds the CD3ε subunit of the TCR. This allows CD8^+^ T-cells to engage target cells that present the GP100 peptide. The cells were added to mOrange expressing A375 tumour cells that were either pre-loaded with GP100 peptide (stimulation condition) or left unloaded (non-stimulation). The target-cell killing assay was performed for 16 hours and killing dynamics was assessed by quantifying the red fluorescence intensity of the target cells. We observed that cells over-expressing FOSL1 produced faster cell killing but only upon engagement of the TCR (Fig. 4H). At the end of the assay, the T cells over-expressing FOSL1 expressed more CD25 than the parental cells, indicating that they were more strongly activated (Suppl. Fig. 5).

## Discussion

We have presented a resource and pipeline for identifying the role of TFs in establishing T-cell states in the setting of preparing cells for T-cell based ACT. We show how bulk RNA-seq can be used to get a deep understanding of how uniformly stimulated cells undergo state transitions and TF changes during both activation and ‘rest periods’ ex vivo. By combining bulk RNA-seq data across a 17-day activation time course with existing single-cell RNA-seq data, using CellOracle, we provide new insights into transcription factor activity in the setting of ACT and identify new opportunities for improving T-cell based therapies, e.g., by using heterologous TF expression to enhance the “super engager” phenotype of CD8^+^ T cells (Dekkers et al, 2023).

Our 17-day time course with anti-CD3/CD28 beads (7-days activation and 10-days rest) leads to a well-defined trajectory of T-cell state formation, matching physiological states from naïve to effector memory, while also allowing us to define the time-points for ex vivo expansion that best produce a given T-cell state. Given such trajectories, additional signals or activation regimens could be used to navigate particular states (i.e., exhaustion, central memory, Tregs). This has important implications for expanding T cells for ACT therapy and research. While a simple bead-based approach is ineffective at generating memory populations known to provide longer-lasting protection and more effective therapeutic outcomes (i.e., central memory T-cells), it is effective at generating effector memory states, which provide for better homing and effector functions (Gattinoni et al, 2011; Klebanoff et al, 2011; Busch et al, 2016; Klebanoff et al, 2012). Our approach is well placed to understand how to produce such states.

We find that the genetic rewiring of T cells is initiated as early as 3 h post-stimulation -a time-point that is often absent in many single-cell datasets. The importance of studying early timepoints was recently exemplified by the finding that T-cell dysfunction/exhaustion is established within hours of tumour encounter (Rudloff et al, 2023), although we did not observe this at our early time point. The lack of an exhaustion signature here may be explained by the use of naïve CD8^+^ T-cells as a starting population, which is known to lead to better outcomes for ACT (López-Cantillo et al, 2022). Additional (i.e., microenvironmental) signals may also be required to drive the early exhausted state seen by Rudloff et al. Examining the later stages in the activation time-course, we find that the expression profile generated during the late resting stage (i.e., 7 days following removal of the stimulus), is profoundly different to that immediately following the proliferative stage. In particular, many classical ‘exhaustion markers’ such as *PDCD1, LAG3, TIM3, TOX2* are highly expressed at Day 7 post-stimulation but we find that their expression is down regulated very rapidly after the stimulus is removed, suggesting that the cells produced by the end of the initial proliferation stage are not truly exhausted. This is an important consideration as many in vitro exhaustion protocols utilise extended bead-based methods to generate proxy-exhausted cells without resting the T cells. As a result, these studies may only be analysing T cells that only transiently exhibit some of the hallmarks of exhaustion (i.e., exhaustion markers) and are not truly exhausted (Jenkins et al, 2023). Our activation time course may help to optimise the point at which ACT will be most effective. For example, the transfer of T cells rested for 2 days rather than 7 days could be counterproductive, owing to the cell’s residual expression of ‘exhausted-like’ gene signatures and functionality.

The regulation of AP1 in T cells is a dynamic and complex process, with the classic AP-1 c-Fos/c-Jun heterodimer driving expression of activation genes including IL-2, and complexes of AP-1 and IRF family members inhibiting c-Jun activity and potentiating exhaustion-related genes. The role of FOSL1 in T cells has not been studied at length, with most insights into its function coming from genome-wide screens. In a genome-wide CRISPR activation screen to identify regulators of interferon-γ production by CD8^+^ T-cells, FOS family members (FOSL1 and FOSL2) were identified as positive regulator hits (Schmidt et al, 2022). In cancer cells, FOSL1 is known to be important for proliferation, apoptosis, and differentiation but what specific role it plays in immune cell subsets is not entirely clear. We showed that FOSL1 expression rapidly increases upon stimulation of naïve T-cells and then its expression declines. This pattern is common to early immediate genes, which comprise the initial response to activation. FOSL1 over-expressing cells had enriched transcriptomic signatures for “super-engager” cells (Dekkers et al, 2023), which correlated with improved killing of target cells. Importantly, in the killing assay, we show that FOSL1 over-expressing cells are not cytotoxic without engagement of the TCR, offering reassurance that FOSL1 would not engender non-specific hyperactivity.

Lastly, our gene/TF-expression trajectories might serve as a resource to help researchers prioritise genes/TFs related to a given function/behaviour when studying T cells at different timepoints during T-cell activation. For example, it could be helpful to prioritise genes/TFs in activated T cells that are experimentally validated to be upregulated in activated versus rested cells as these are more likely to be essential to the T-cell function of interest at that stage in activation. Our dataset can also serve as a reference to identify a transitory or stationary T-cell state and infer the directionality of T-cell state transitions. This may be of use when it’s possible only to analyse a single ‘snapshot’ of the T-cell state. At a more systems-level, the time course RNA-seq dataset generated in this study can serve as a guide to understanding when the key drivers of signalling occur at a gene expression level and to refine experimental designs for ‘-omics’ datasets.

## Limitations

Here we have used only naïve CD8^+^ T-cells, a bead-based activation approach, and T-cell states and transitions driven by signals 1 and 2 (i.e., TCR and costimulatory signals, respectively). Although signals 1 and 2 are critical for driving T-cell activation, cytokines and different environmental signals will undoubtedly influence the T-cell state trajectory and the expression patterns of TFs. However, our goal was to first establish a pipeline to examine TF networks driven by the core activating receptors of T cells, and we are now set to expand our approach to examine different starting populations and methods of activation in future experiments.

## Materials and methods

### Cell culture

Previously established T-cell stimulating (TCS) cell line expressing an anti-human CD3 single-chain fragment fused to human CD14 were utilised (Leitner et al, 2010). The Jurkat T cells and TCS cells were cultured in RPMI 1640 medium (Gibco) supplemented with 10% (v/v) fetal bovine serum (FBS), 1% (v/v) Penicillin/Streptomycin/Neomycin (final concentrations of 50 U/ml Penicillin, 50 μg/ml Streptomycin, 100 μg/ml Neomycin), 1% (v/v) glutamine and 1% (v/v) HEPES (collectively referred to as ‘complete-RPMI’). HEK-293T cells for lentiviral transfection were cultured in DMEM (Gibco), with additions of the same supplements as above. CD8+ primary T cells were cultured in complete RPMI supplemented with/without 50U/ml of IL-2 (Peprotech). In primary cell experiments, cells were cultured in media without IL-2 for 24 hours prior to assay. All cells are incubated at 37°C with 5% CO2.

### Activation of naïve cells and RNA extraction

Naïve CD8+ T cells were isolated from PBMCs using Naïve CD8+ T Cell Isolation Kit from Miltenyi (catalog number: 130-093-244) using the manufacturer’s instructions. Approximately 1 million cells were used for activation using Dynabeads™ (anti-CD3/anti-CD28 beads) from ThermoFisher (catalog number: 11131D). The bead to cell ratio was 3:1. RNA was extracted from naïve or activated CD8+ T cells using RNeasy Kit (Qiagen, catalog number: 74104) following manufacturer’s instructions.

### Cell culture and transductions

Jurkat T cells were seeded at one million cells/well in 6-well plates, and 1ml of lentiviral supernatant was added to each well. Transduction efficiency was measured via flow cytometry and validated via western blot. FOSL1 (NM_005438.5) expression construct was purchased from Genescript and the insert was cloned into pHR lentiviral vector.

### 2 Cell Assay (2CA)

The two cell assay was performed as previously described. Briefly, Jurkat T cells were co-cultured with TCS in a 2:1 ratio. To generate different stimulation conditions, Ultra-LEAF anti-human CD86 and PD-L1 antibodies (Biolegend (Clone #IT2.2 and #29E.2A3 respectively)) were added to final concentration of 1 μg/ml. Cells were harvested after 24 hours. TCS cells were stained with PE-conjugated anti-mouse CD45 antibody (Clone #I3/2.3; BioLegend), whereas Jurkat or primary cells were stained for CD69 activation marker (Clone #FN50), followed by flow cytometric analysis. Flow cytometry was performed on an LSRFortessa using FACSDiva software. Data was analyzed using the CytoExporeR package in R. All analyses were gated on viable and single cells, which was determined according to their FCS/SCC profile.

### Activation, negative TCS selection of Jurkat T cells

Jurkat T cells were seeded at 1 million cells/well in 6-well plates and were either activated with an equal number of TCS cells, or not activated. Cells were harvested after 24 hours, and all samples were treated to negatively isolated TCS cells. A final concentration of 0.5 μg/ml biotin-conjugated mCD45 antibody (clone #30-F11, BioLegend: 103103) was added to the cells on ice. After 15 minutes, the cells were washed with MojoSort buffer (5% (w/v) BSA, 2 mM EDTA in PBS at pH 7.2, filtered with 0.22 μm SFCA syringe filter before use), and incubated with 2 μl MojoSort Streptavidin Nanobeads (BioLegend catalog # 480015) on ice for 15 minutes. CD45 labelled TCS cells were separated using a magnetic rack, and Jurkat cells were collected into a new tube. Purity and activation status of Jurkat T cells in the final mixture was validated via flow cytometry. RNA was extracted from Jurkat cells using the RNeasy kit (Qiagen) using manufacturer’s instructions.

### Transduction of primary cells

Primary CD8+ T cells were isolated using RosetteSep™ Human CD8+ T Cell Enrichment Cocktail (Stemcell, Catalog number 15063) using manufacturer’s instructions. Cells were activated using Human T-Activator CD3/CD28 ImmunoCult™ (Stemcell, Catalog number 10971) 2 days prior to transduction. Recombinant Human Fibronectin Fragment (RetroNectin, TakaraBio, Catalog number T100A) reagent was used to aid co-localisation of target cells and virions. 300 μl of 25 μg/mL RetroNectin diluted in sterile PBS was applied to Non-Tissue Culture Treated 24-well plates at least 24 hours prior to transduction at 4 °C, or 2 hours at room temperature. To bind virus particles, wells were blocked with 2 % BSA for 30 minutes before centrifugation at 2000g at 32 °C for 90 minutes with 1.5 mL of viral supernatant. To bind target cells, 1.4ml viral supernatant was aspirated, and 1 million activated CD8+ T cells were immediately added to each well. The plate was spun at 526g at 32°C for 2 minutes. Transduced cells expressing GFP were isolated using FACS sorter 3 days post-transduction and used in subsequent assays.

### Killing Assay using effector T-cells

FOSL1 over-expressing CD8+ T cells were generated using lentiviral transduction. 48-hours post-transductions, cells expressing GFP were sorted and cultured for a further 24 hours. Melanoma cell line, A375, engineered to express mOrange were loaded with GP100 peptide for ∼4 hours. Untransduced CD8+ T-cells and CD8+ T-cells over-expressing FOSL1 cells were added into target cells together with bispecific antibody (10^−8^ M) and the plate was imaged every 2 hours using the Incucyte Live-Cell Analysis System. After 16 hours, cells were harvested and stained using anti-CD25 antibody, measured using flow cytometry. Killing levels were quantified from the Incucyte by measuring the area and the intensity of ‘red’ fluorescence.

### Bulk RNA-seq analysis

FASTQ files were aligned using STAR version 2.7.3a and quantified using featureCounts version 2.0.6. Ensembl Homo sapiens GRCh38.86 was used as the reference genome. Lowly expressed genes were filtered out, and gene expression distributions of each sample were normalised using edgeR version 3.42.2. Differential expression analysis was performed using limma version 3.54.2. Genes with absolute log-fold-change>1.5 and adjusted p.value<0.05 were defined as significant.

### Construction of network using GENIE

Raw count files from featureCounts were filtered using the rowSums and cpm functions to retain genes with a CPM of >1 in at least one sample, normalised using the calcNormFactors function, and log-transformed using the cpm function in edgeR version 3.40.2. Taking the log-CPM values as input, the top 35% most highly variable genes (HVGs) were identified using the getTopHVGs function in scran version 1.26.2. The raw count values of the HVGs were fed into GENIE3 version 1.20.0 to build regulatory networks. Only transcription factors (TFs) reported in Lambert et al., 2018 were used as candidate regulators (Lambert et al, 2018). The software yielded a list of regulatory links, each of which contains a regulator, a target gene and a weight of the link. A threshold of 0.04 was applied to filter out less significant links. The remaining links were then imported into Cytoscape, in which networks were built and visualised, providing insights into gene regulation.

### CellOracle implementation

Raw count scRNA-seq files of naïve and memory CD8+ T cells (0, 12, 24 h post-stimulation) from Bibby et al., 2022 were preprocessed with Scanpy version 1.9.3 and then analyzed with CellOracle version 0.12.0 (Bibby et al, 2022; Kamimoto et al, 2023). Cells were filtered to retain those with gene counts 200-7,500 (naïve) or 200-6,000 (memory), <15% mitochondrial counts and >65 (out of 97) housekeeping genes compiled in Tirosh et al. (Tirosh et al, 2016). Following normalization of each cell by total counts of all genes, and log transformation, HVGs were identified using the highly_variable_genes function in Scanpy. Scanorama version 1.7.3 was then applied to remove batch effects and integrate datasets across time points. Thirty principal components were used to compute a neighbourhood graph of cells, and the Louvain algorithm with a resolution of 0.6 (naïve) or 0.4 (memory) was used to cluster the neighbourhood graph. In the naïve T-cell dataset, two disjoint clusters, naïve and TEMRA, were identified by the algorithm, and only the naïve subset was used in the CellOracle analysis. Similarly, the memory T-cell dataset exhibited two disjoint clusters, CD8αα and CD8αβ, and only the CD8αβ subset was used. To infer the progression of these cells, the DPT algorithm was used to calculate pseudotime. CellOracle GRN models were constructed using the pre-built promoter base GRN for human (hg19) as the reference genome. The models simulated changes in cell identity in response to TF perturbations. To properly interpret the simulation results, a vector recapitulating the developmental flow of the CD8+ T cells, derived from DPT pseudotime analysis, was built. Perturbation scores (PS) were then calculated by comparing this vector with the simulated perturbation vector. Positive PS suggest that the TF perturbation would promote differentiation, whereas negative PS block differentiation. To ensure that only TFs were selected, only TFs that occur in both the base GRN and Lambert et al., 2018 were used in downstream analysis (Lambert et al, 2018).

### Data integration with publicly available RNA-seq dataset

FASTQ files generated in Project PRJNA744261 were downloaded from the European Nucleotide Archive (ENA). The files were aligned using STAR version 2.7.3a and quantified using featureCounts version 2.0.6. Ensembl Homo sapiens GRCh38.86 was used as the reference genome. Lowly expressed genes were filtered out using the filterByExpr function, and gene expression distributions of each sample were normalised using the calcNormFactors function in edgeR version 3.42.2. The Combat function in sva version 3.48.0 was used to adjust for batch effects in our datasets and the publicly available datasets.

## Data Availability

All the raw sequencing data together with the associated metadata generated from this work has been deposited to GEO (GSE247647).

## Acknowledgements

We would like to thank Mai Voung for helping with cloning and Evangelia Petsalaki for critical reading of the manuscript and for providing helpful comments.

## Declarations

YL, EJ, AMS, and SS are named as inventors on a patent application describing the methods and results included in this publication.

## Author contribution

Conceptualization: Sumana Sharma, Edward Jenkins, Ana Mafalda Santos, Simon Davis

Methodology: Yuan Lui, Sumana Sharma, Joseph Clarke, Mateusz Kotowski, Sydney Mullin, Ana Mafalda Santos

Software: Sumana Sharma, Yuan Lui

Validation: Emily Ng, Sumana Sharma

Formal analysis: Sumana Sharma, Yuan Lui

Investigation: Edward Jenkins, Ana Mafalda Santos, Sumana Sharma, Emily Ng

Resources: Simon Davis

Data curation: Yuan Lui

Writing - Original Draft: Sumana Sharma, Edward Jenkins

Writing - Review & Editing: Sumana Sharma, Edward Jenkins, Emily Ng, Ana Mafalda Santos, Simon Davis

Visualization: Sumana Sharma, Yuan Lui

Supervision: Sumana Sharma, Simon Davis

Project administration: Sumana Sharma, Ana Mafalda Santos

Funding acquisition: Simon Davis

