## Supplementary Figures for "Cell state and transcription factor modulation during extended ex vivo CD8^+^ T-cell expansion"

**[1]MRC Human Immunology Unit, John Radcliffe Hospital, University of Oxford, Oxford, United Kingdom**

***authors contributed equally**

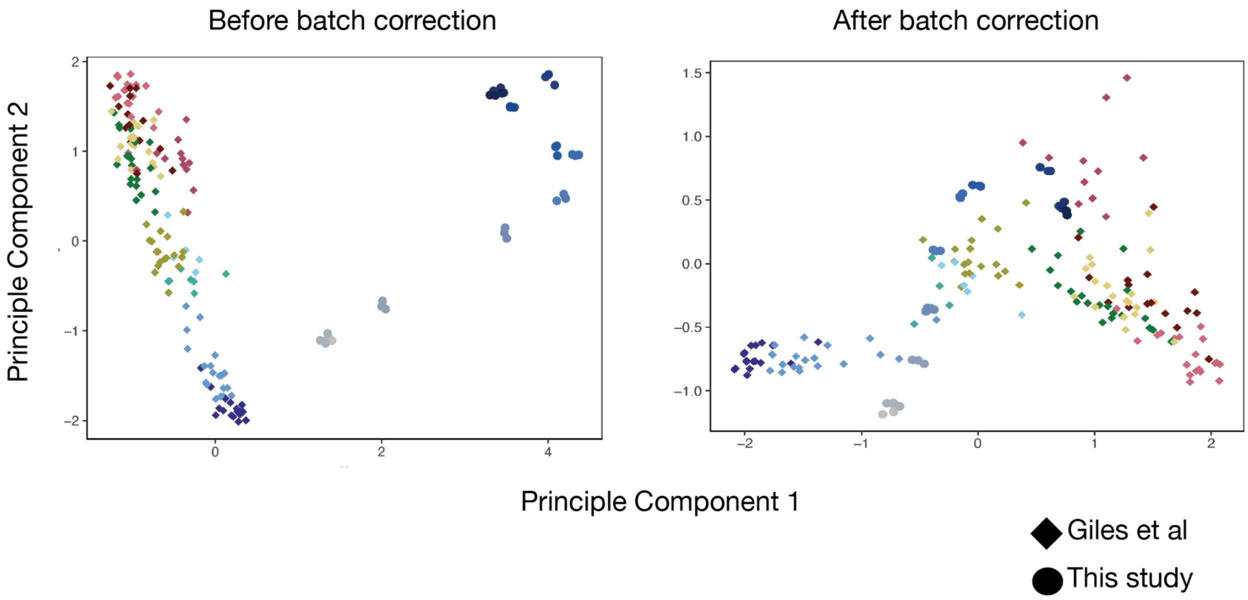


**Supplementary Figure 1**: PCA of CD8+ T cells from this study and from Giles et al. before and after batch correction using ComBAT. No genes were removed during the batch correction.

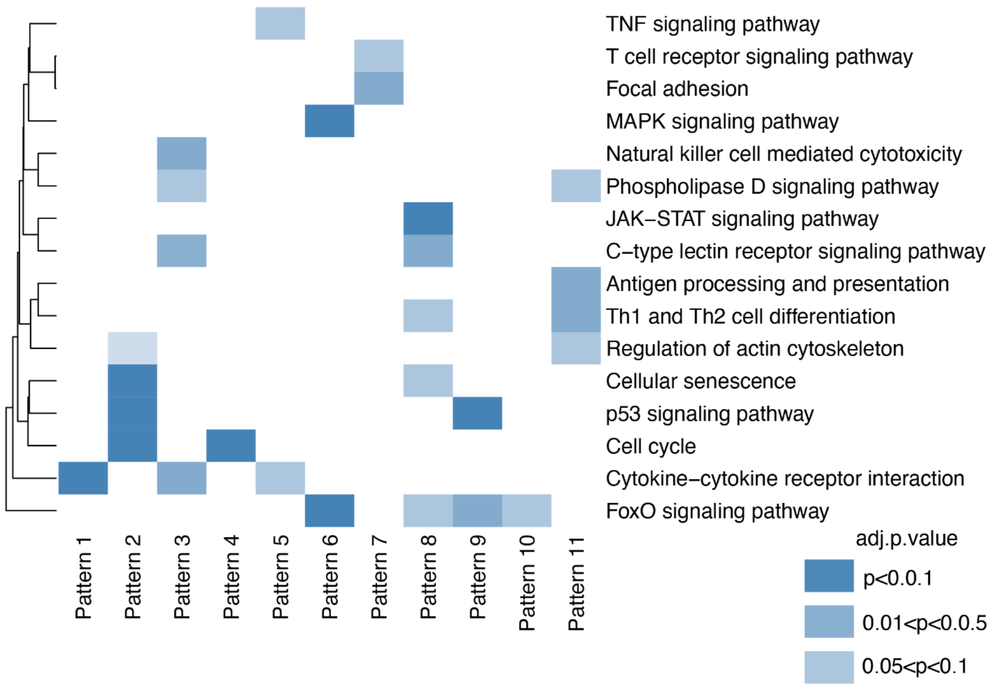


**Supplementary Figure 2:** Enriched pathways in each pattern identified using enrichr.

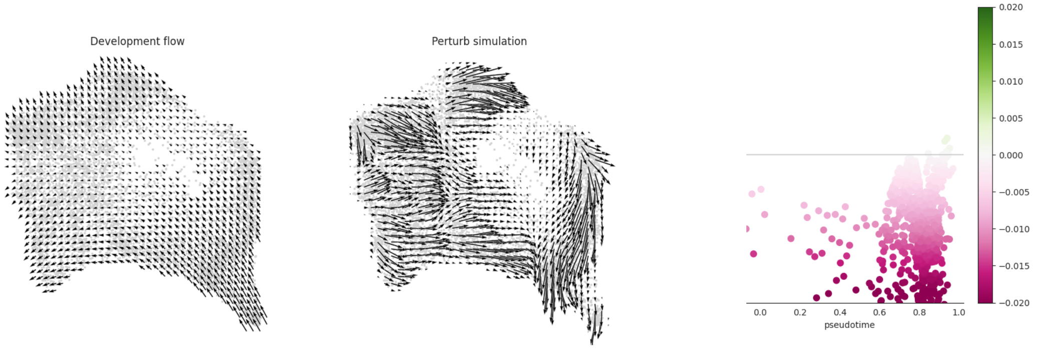


**Supplementary Figure 3:** In silico KO of *NFKB1* in memory cells using CellOracle. The developmental flow of memory cells (left panel) is perturbed by KO of *NFKB1 (*middle panel). The negative PS score (right panel) implies that TF KO delays or blocks the transition to an activated/proliferative state.

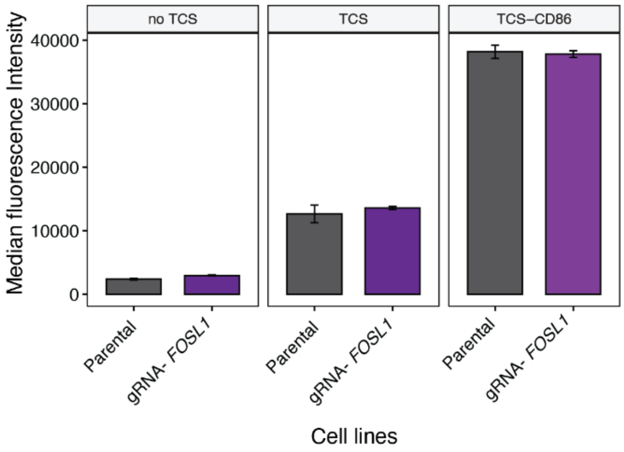


**Supplementary Figure 4**: Targeting FOSL1 in Jurkat cells does not affect signalling outcome. FOSL1 in Jurkat cells was targeted using sgRNA. Parental and KO cells both responded to the same levels post co-culture with TCS as indicated by the expression of CD69.

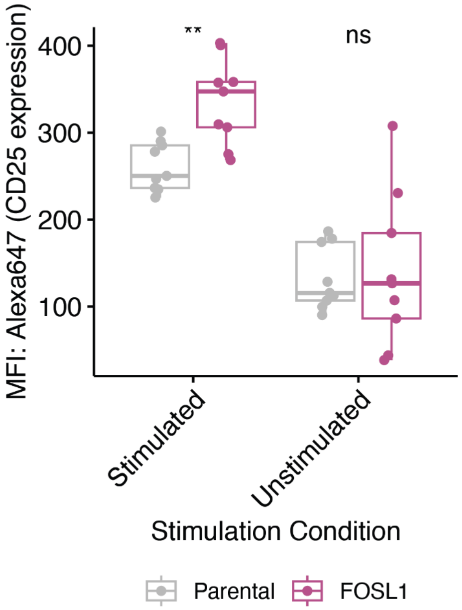


**Supplementary Figure 5:** CD25 expression on parental and FOSL1 over-expressing cells post tumour killing.
